# iPSC-derived Vascular Smooth Muscle Cells in a Fibronectin Functionalized Collagen Scaffold Augment Endothelial Cell Morphogenesis

**DOI:** 10.1101/2021.07.22.453420

**Authors:** Kaiti Duan, Biraja C. Dash, Daniel Sasson, Henry C. Hsia

**Affiliations:** Section of Plastic Surgery, Department of Surgery Yale School of Medicine, Yale University, New Haven, USA; Department of Biomedical Engineering, Yale University, New Haven, USA

**Keywords:** Induced pluripotent stem cell, vascular smooth muscle cells, fibronectin, vascular network formation, angiogenesis, integrin signaling

## Abstract

Tissue-engineered constructs have immense potential as autologous grafts for wound healing. Despite the rapid advancement in fabrication technology, the major limitation is controlling angiogenesis within these constructs to form a vascular network. Here, we aimed to develop a 3D scaffold that can regulate angiogenesis. We tested the effect of fibronectin and vascular smooth muscle cells derived from human induced pluripotent stem cells (hiPSC-VSMC) on the morphogenesis of endothelial cells. The results demonstrate that fibronectin increases the number of endothelial networks. However, hiPSC-VSMC in the presence of fibronectin further substantiated the number and size of endothelial networks. A mechanistic study shows that blocking αvβ3 integrin signaling between hiPSC-VSMC and fibronectin impacts the endothelial network formation. Collectively, this study set forth initial design criteria in developing an improved pre-vascularized construct.

## Introduction

Tissue-engineered constructs provide an alternative to autologous grafts for the treatment of non-healing wounds [1]. However, the constructs fail to stably engraft in chronic wounds with vascular insufficiency requiring additional interventions [2]. The primary reason for this failure is the lack of vascularization in the implanted tissue-engineered constructs and cell death [3]. A pre-vascularized construct improves engraftment and survival after transplantation and helps in establishing a functional vasculature in the host [4]. In sum, this evidence suggests that pre-vascularized scaffolds can promote chronic wound healing.

Morphogenesis of endothelial cells is a prerequisite for vessel formation [5]. During morphogenesis, the ECs proliferate, migrate and self-assemble to form a tube-like structure [6]. Studies reveal that extracellular matrix (ECM) such as fibronectin regulate vascular remodeling via integrin signaling. Gerecht and colleagues in one such study demonstrated the role of fibronectin in matrix assembly and EC morphogenesis. They studied the role of polymerized fibronectin on vascular tubulogenesis on a decellularized matrix. Most importantly, they showed the importance of fibronectin and integrin αvβ3 and α5β1 in EC morphogenesis [7].

One of the vascular tissue engineering strategies is to utilize stromal cells to rebuild patent vessels in a scaffold matrix for subsequent transplantation [8]. The generation of vascular networks containing vascular smooth muscle cells (VSMC) is highly relevant as they play an integral part in forming vasculatures both in a natural and pathological process [9]. In more recent works, we and others have demonstrated the derivation of VSMC from human-induced pluripotent stem cells (hiPSC) [10, 11]. This hiPSC-VSMC is pro-angiogenic and promotes wound healing via paracrine secretion [11]. Moreover, hiPSC is considered less immunogenic and provides a source of rejuvenated cells with no ethical concerns, further increasing their attractiveness as a source for VSMCs for chronic wound healing [12].

In our recent studies, we showed that changing the three-dimensional environment of the scaffolds, whether it is to increase the collagen density or addition of fibronectin, promotes hiPSC-VSMC’s pro-angiogenic functions via paracrine secretions [11, 13]. Engineering improved pre-vascularized constructs with the support of a primed hiPSC-VSMC in a deliverable ECM-based scaffold offer unmet opportunities to treat non-healing wounds. In the current study, we intended to investigate the effects of pro-angiogenic hiPSC-VSMC on endothelial morphogenesis. We hypothesized that pro-angiogenic hiPSC-VSMC provides essential cues for the growth and maintenance of EC-based networks in a fibronectin functionalized collagen scaffold. This study is a critical step toward the fabrication of pre-vascularized tissues.

## Methods

### Cell culture

Human iPSC-VSMC were differentiated using a previously established protocol [14]. hiPSCs were cultured in feeder-free conditions for 4 days, and then cellular colonies were collected after treating with Dispase for 15 minutes at 37°C. The released cellular colonies were transferred to a low attachment plate to form embryoid bodies (EB) and treated in an mTESR medium. After 24 hours, the culturing medium was changed to a mixture of mTESR and a customized EB differentiation medium (DMEM high glucose + 10% FBS + 1% non-essential amino acid (v/v) + 2mM L-glutamine and 0.012 mM 2-mercaptoethanol) at a 1 to 3 ratio. After 72 hours, the medium was changed to the customized EB differentiation medium only. The EBs were cultured for another 48 hours and transferred to a gelatin-coated plated and was further cultured for another 5 days followed by culturing them on a Matrigel-coated plate for a week. The resulting differentiated cells were cultured in a commercially available smooth muscle cell medium (SmGM-2) and verified with immunofluorescence staining of SM-22α and calponin. After verification, iPSC-VSMCs were either stored in liquid nitrogen or plated onto gelatin-coated Petri dishes.

### HUVEC culture

Primary human umbilical vein endothelial cells (HUVECs) were cultured in Endothelial Cell Growth Medium (EGM)-2, which was replenished every 48 hours, until at least 80% confluence. All HUVECs used in our experiments were below passage number 7.

### Fibronectin collagen scaffold production

The fibronectin functionalized collagen scaffolds (collagen+fibronectin) were produced in the following steps [13]. The initial concentration of rat tail type-1 collagen used was 5mg/ml. To maintain the integrity of the materials, the entire scaffold formation process was done on ice. To achieve an overall density of 4mg/ml collagen, the following ingredients were mixed within an Eppendorf tube in the following order: 400 μl of type-I collagen, 50 μl of MEM, then 8.4 μl of 1M NaOH and 50 μl of 1x PBS or 1mg/ml of fibronectin. The formulations of each scaffold group are listed in Table 1. The mixture was gently mixed to prevent bubble formations. After the mixture color change from yellow to a light purple tint, respective cells, HUVEC, or a combination of hiPSC-VSMC and HUVEC of 1:4 ratio were added. HUVEC only scaffolds contained about 1×106 cells, and combination scaffolds contained 2×105 hiPSC-VSMC and 8×105 HUVEC. The cell-scaffold mixture was then further gently mixed and distributed as 100μl scaffold aliquots into 96 well cell culture plates containing approximately 2×105 of total cells. The plates were then incubated in the cell culture incubator for at least 30 minutes at 37°C for the gelation process. The following table contained the appropriate amount of components for each scaffold mixture. After the scaffolds achieved gelation, 200μl of EGM-2 or combination media (1:1 of EGM-s and SmGM-2) were added on top of the scaffolds and were incubated for 7 days, while changing media every 72 hours. The various groups were HUVECs and coculture of HUVECs and hiPSC-VSMCs in collagen scaffolds (Col-I), and collagen+fibronectin scaffolds (Col-I+Fib).

**Table 1:** Various formulations of collagen+fibronectin scaffolds with hiPSC-VSMCs and HUVECs in a total volume of 500 μl.

| Samples | Density (mg/ml) | Collagen (µl) | 10x MEM (µl) | 10x PBS (µl) | 1M NaOH (µl) | Fibronectin 1mg/ml (µl) | hiPSC VSMC $8 \times 10^3/\mu\text{l}$ | HUVEC $16 \times 10^3/\mu\text{l}$ |
| --- | --- | --- | --- | --- | --- | --- | --- | --- |
| Col-I+EC | 4 | 400 | 50 | 50 | 8.4 | 0 | 0 | 62.5 |
| Col-I+Fib+EC | 4 | 400 | 50 | 0 | 8.9 | 50 | 0 | 62.5 |
| Col-I+hiPSC-VSMC+EC | 4 | 400 | 50 | 50 | 8.4 | 0 | 25 | 50 |
| Col-I+Fib+hiPSC-VSMC+EC | 4 | 400 | 50 | 0 | 8.9 | 50 | 25 | 50 |

### αvβ3 integrin signaling inhibition experiment

Another set of fibronectin functionalized collagen scaffolds containing either HUVECs or a combination of hiPSC-VSMCs and HUVECs were constructed under the same condition as described above. 24 hours after the initial culture, 2 μl echistatin (100nM), an inhibitor of αvβ3, was added into the culturing medium and was refreshed every 2 days. At the end of the 7 days, all the scaffolds were harvested and fixed with 4% paraformaldehyde for immunofluorescence staining.

### Immunofluorescence staining

After day 7 the scaffolds were harvested and washed with 1xPBS and then fixed with 4% paraformaldehyde for at least 10 minutes. Then the scaffolds were washed 3 times with 1xPBS for 10 minutes and blocked with 5% Bovine Serum Albumin in PBS and 0.05% Tween20 (BSA-PBST) for 1 hour at room temperature. The scaffolds were incubated with CD-144 and SM-22-α primary antibodies at 1:200 dilution overnight at 4°C. On the next day, the scaffolds were washed three times with PBST (tween20 0.05%), and then the samples were incubated with secondary antibodies tagged with Alexa Fluor® 488 for 1 hour at room temperature. Dapi was used as a counterstain at 1:1000 dilution for 10 minutes. The scaffolds were then washed, mounted on slides, and images were taken under a confocal microscope.

### Endothelial cell network quantification

Images from each scaffold were taken from 5 different microscopic fields independently by two different lab members and then the number of vascular networks was counted by one individual. The individual also measured and separated the sizes of the networks with cutoff boundary at 50 μm in length. The sizes of the vascular networks (> 50 μm and < 50 μm) were determined by measuring the shortest side of each vascular network.

## Results

### Fibronectin increased the number of endothelial cell networks

The CD-144 staining of the HUVEC only scaffolds demonstrate the formation of EC networks over seven days (Figures 1 and 2). The fibronectin functionalized group shows an enhanced number of EC networks than the control collagen scaffolds (Figure 1B and 2A). We see a modest increase in the number of networks from an average of 8 to 11. However, we do not see any difference between the number of different sizes (> 50 μm and < 50 μm) in both collagen only and fibronectin functionalized scaffolds (Figure 2B and C). Treatment with αvβ3 inhibitor reduced the number of EC networks in the fibronectin functionalized groups close to that of the control collagen scaffolds (Figure 1B and 2A). However, we do not observe a change in the size of EC networks (Figure 2B and C).

**Figure 1:**
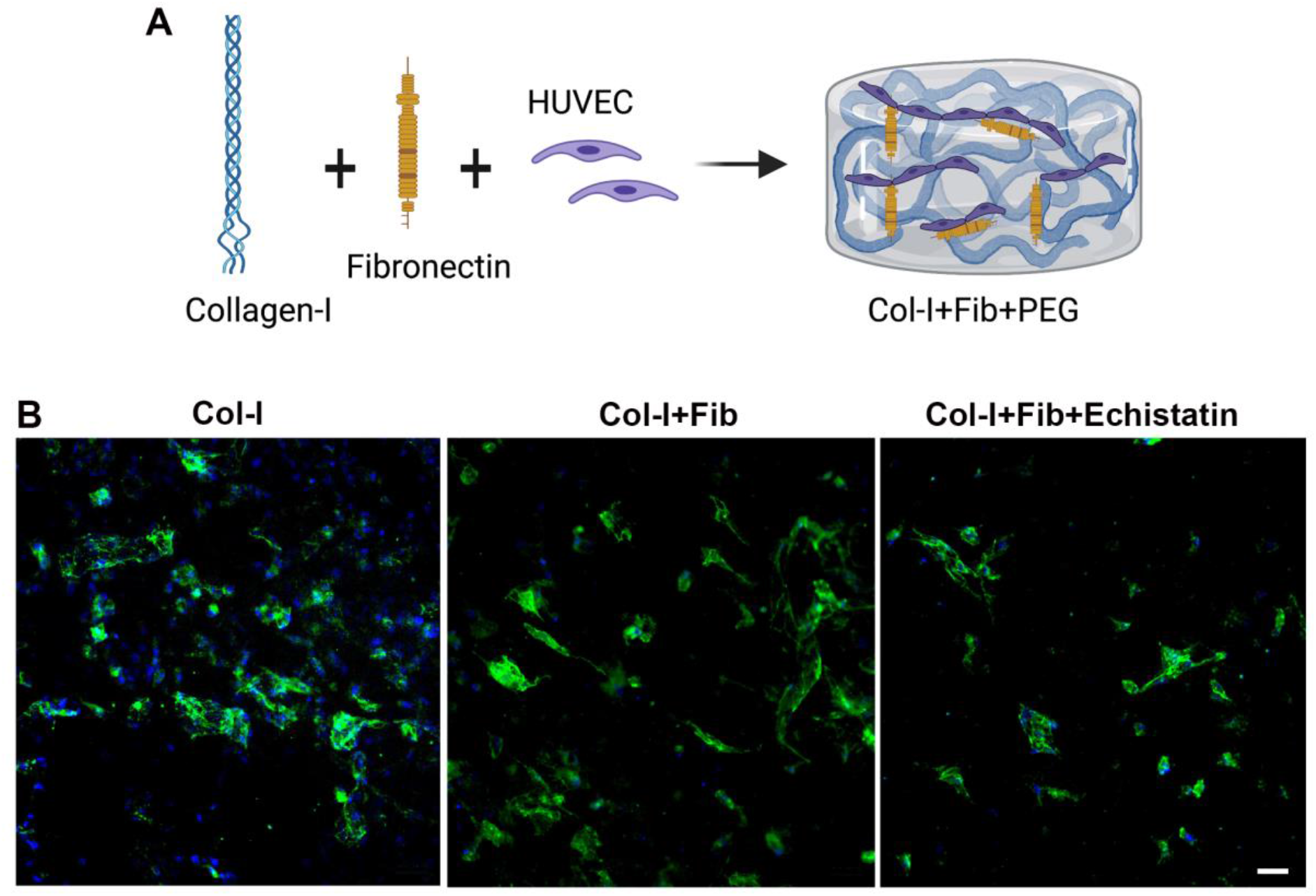
Characterization of HUVEC only scaffolds. (A) The schematic shows HUVEC in fibronectin functionalized collagen scaffold forming EC networks. (B) Fluorescent images of HUVEC-only scaffolds stained with CD144 on Day 7. The different scaffolds were collagen only (Col-I), fibronectin functionalized collagen scaffold (Col-I+Fib), and Col-I+Fib scaffold treated with Echistatin (Col-I+Fib+Echistatin). The scale bar measures 50 μm.

**Figure 2:**
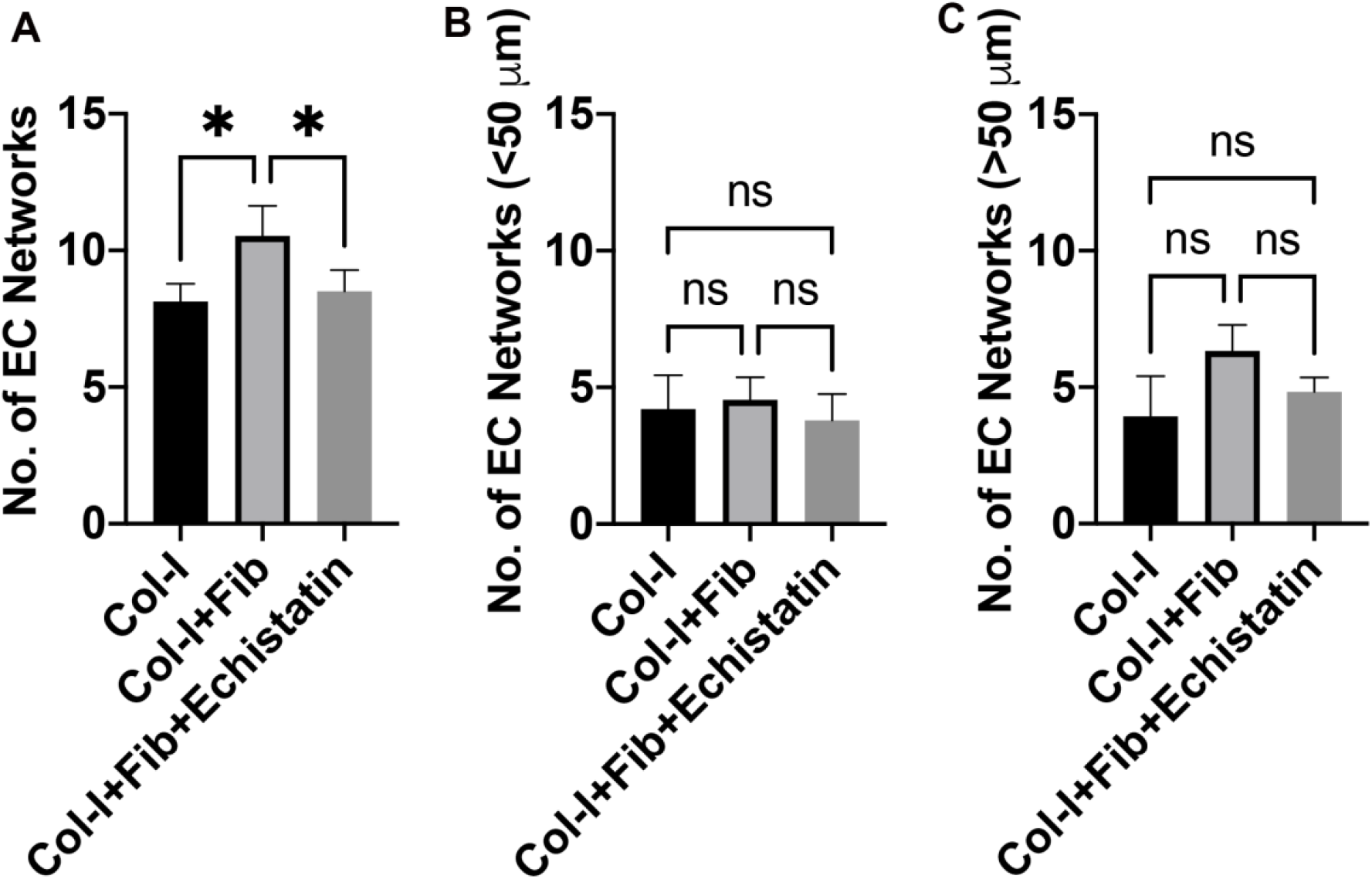
Quantification of EC networks in HUVEC only scaffold. The graphs show quantification of (A) total number of EC networks, (B) networks less than 50 μm, and (C) networks more than 50 μm. The different scaffolds were collagen only (Col-I), fibronectin functionalized collagen scaffold (Col-I+Fib), and Col-I+Fib scaffold treated with Echistatin (Col-I+Fib+Echistatin). Statistical significance was determined using One-way ANOVA (* p< 0.05, n=3).

### hiPSC-VSMC in the presence of fibronectin enhanced the number and size of endothelial cell networks

We quantified the number of CD144 stained EC networks in the co-culture of hiPSC-VSMC and HUVEC in the fibronectin functionalized and control scaffolds (Figure 3 and 4). We see a similar trend as HUVEC-only culture in the fibronectin functionalized scaffold compared to control (Figure 3B and 4A). However, the co-culture of hiPSC-VSMC and HUVEC shows a substantial increase in the number of EC networks from an average of 8 to 16 (Figure 4A). We do not observe any significant difference in the number of smaller EC networks (< 50 μm) between fibronectin functionalized and collagen scaffolds (Figure 4B). At the same time, we see nearly 78% of the total EC networks are larger than 50 μm in size in the fibronectin group compared to 54% in the case of control collagen scaffolds (Figure 4C). The number of > 50 μm EC networks is three times that of control collagen scaffolds (Figure 4C). Treatment with αvβ3 inhibitor reduced the EC networks in the fibronectin functionalized groups by close to 20% and affected their size (Figure 4A, 3B, 4B, and 4C).

**Figure 3:**
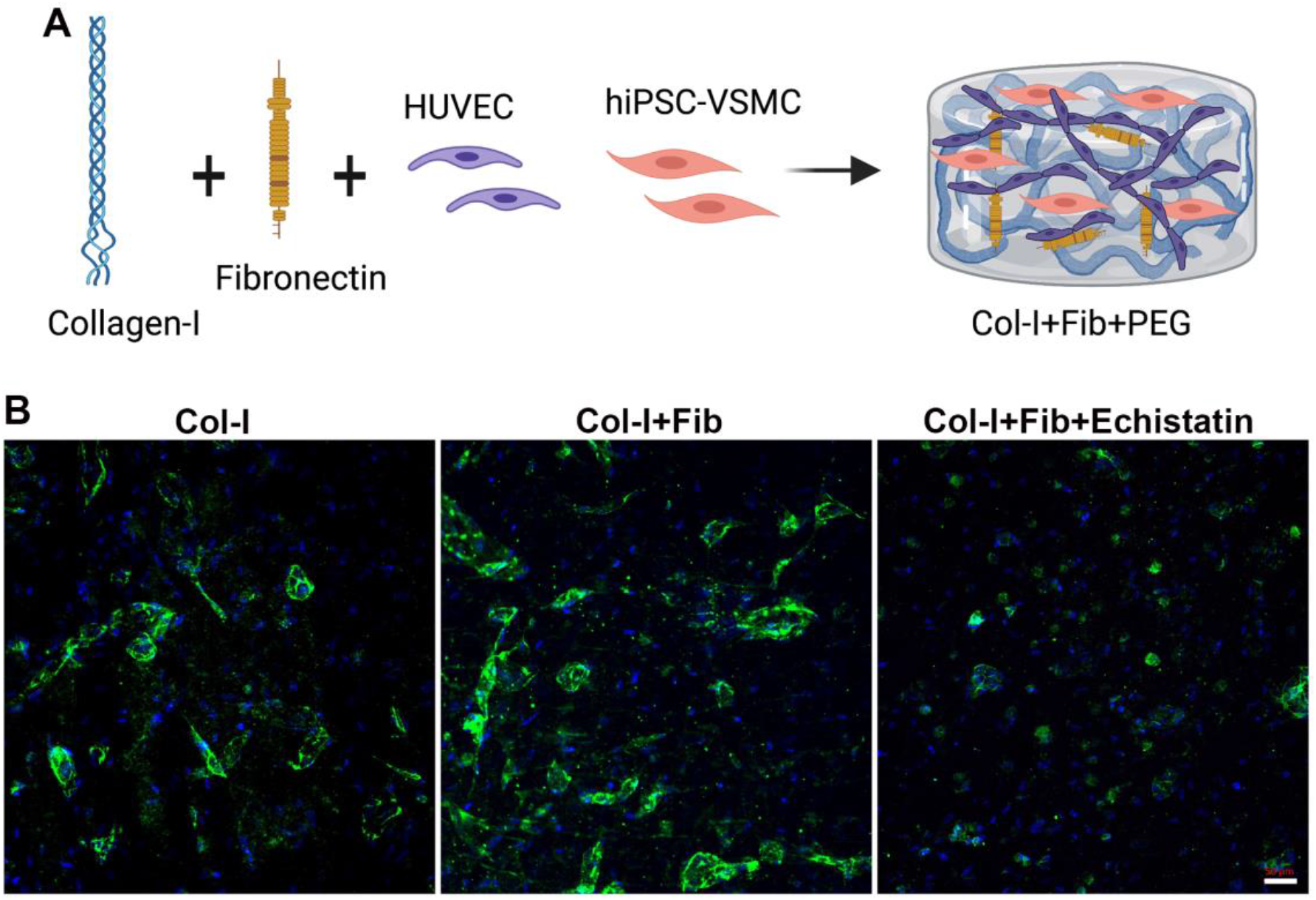
Characterization of hiPSC-VSMC and HUVEC scaffolds. (A) The schematic shows hiPSC-VSMC and HUVEC in fibronectin functionalized collagen scaffold forming EC networks. (B) Fluorescent images of hiPSC-VSMC and HUVEC scaffolds stained with CD144 on Day 7. The different scaffolds were collagen only (Col-I), fibronectin functionalized collagen scaffold (Col-I+Fib), and Col-I+Fib scaffold treated with Echistatin (Col-I+Fib+Echistatin). The scale bar measures 50 μm.

**Figure 4:**
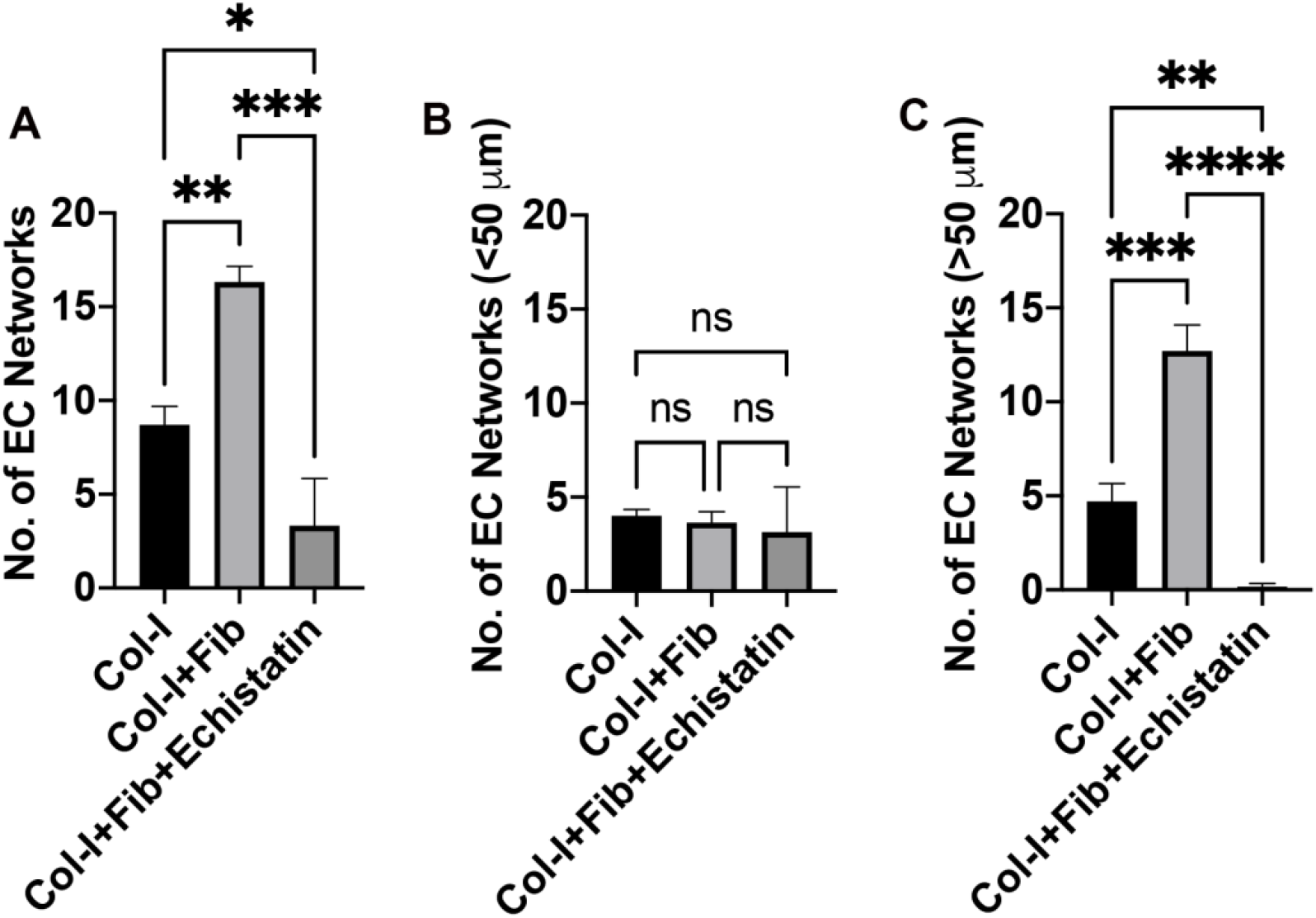
Quantification of EC networks in hiPSC-VSMC and HUVEC scaffolds. The graphs show quantification of (A) total number of EC networks, (B) networks less than 50 μm, and (C) networks more than 50 μm. The different scaffolds were collagen only (Col-I), fibronectin functionalized collagen scaffold (Col-I+Fib), and Col-I+Fib scaffold treated with Echistatin (Col-I+Fib+Echistatin). Statistical significance was determined using One-way ANOVA (* p< 0.05, ** p < 0.01, *** p < 0.001, **** p < 0.0001, n=3).

## Discussion

This study aims at optimizing a proangiogenic scaffold using fibronectin and hiPSC-VSMC. Fibronectin, an ECM protein, regulates the proangiogenic function of EC and maintains their fenestrae structures [7, 15]. In this study, we demonstrated that by interrupting the fibronectin and αvβ3 integrin interaction by echistatin, there is a significant decrease in the overall viability of HUVEC (Supplementary Figure 1). This data indicates that fibronectin interaction with αvβ3 integrin is required to maintain EC viability. In addition, we demonstrate that fibronectin functionalized collagen scaffold promotes endothelial network formation via αvβ3 integrin. This result is consistent with previous reports [7].

We recently demonstrated that fibronectin primes hiPSC-VSMC with enhanced paracrine secretion. Towards the goal of developing a pre-vascularized scaffold, we attempted to use hiPSC-VSMCs in the presence of fibronectin to regulate EC morphogenesis. The primed hiPSC-VSMC in the fibronectin functionalized scaffold significantly increased the quality of EC networks by increasing the size of vascular networks. Interestingly, treatment with the αvβ3 inhibitor profoundly affects the size of EC networks. This observation is a stark departure from the HUVEC-only echistatin treated scaffolds, where we do not observe a difference in the size. We have previously reported that blocking αvβ3 signaling negatively affects the secretion of proangiogenic factors such as bFGF and MMP-2 in hiPSC-VSMC [13]. Collectively, our data indicates that hiPSC-VSMC in the presence of fibronectin actively enhances the morphogenesis of HUVEC in forming large vascular networks. Our study further suggests the importance of fibronectin and hiPSC-VSMC interaction and paracrine secretion in EC morphogenesis. This finding is a first and critical step towards developing an improved pre-vascularized scaffold for wound healing.

## Author Contributions

K.D. and B.C.D. conceived the study. B.C.D. and H.C.H. procured the funding. K.D. and B.C.D designed the experiments. K.D. and D.S. performed the experiments. K.D. and B.C.D wrote the manuscript. All the authors participated in data analysis, discussed the results, and reviewed the manuscript.

## Acknowledgements

The authors would like to thank the core research facilities at the Yale Department of Surgery.

