## Supplemental Information for "iPSC-derived Vascular Smooth Muscle Cells in a Fibronectin Functionalized Collagen Scaffold Augment Endothelial Cell Morphogenesis"

Running Title: hiPSC-VSMC Promotes EC Morphogenesis

### Equal Contribution

Address: Yale University School of Medicine, 330 Cedar St., BB 3rd Floor, PO Box 208041, New Haven, Connecticut 06510, Telephone: 203 737-2049, Fax: 203 785-5714

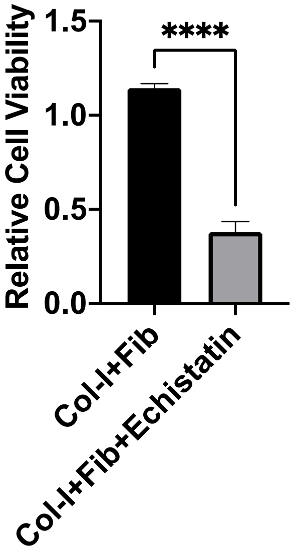


Figure S1: HUVEC viability characterization. The AlamarBlue assays showing relative viability HUVEC on fibronectin functionalized collagen scaffolds (Col-I+Fib) and Col-I+Fib scaffold treated with Echistatin (Col-I+Fib+Echistatin). Statistical significance was determined using One-way ANOVA (**** $p < 0.0001$, n=5).
